# Generative prediction of real-world prevalent SARS-CoV-2 mutation with *in silico* virus evolution

**DOI:** 10.1101/2024.11.28.625962

**Authors:** Xudong Liu, Zhiwei Nie, Haorui Si, Xurui Shen, Yutian Liu, Xiansong Huang, Tianyi Dong, Fan Xu, Zhixiang Ren, Peng Zhou, Jie Chen

**Affiliations:** School of Electronic and Computer Engineering, Peking University, Shenzhen, China; Pengcheng Laboratory, Shenzhen, China; Guangzhou Medical University, Guangzhou, China; State Key Laboratory of Respiratory Disease, Guangzhou, China; Guangzhou National Laboratory, Guangzhou, China; School of Computer Science, Peking University, Beijing, China; State Key Laboratory of Virology and Biosafety, Wuhan Institute of Virology, Chinese Academy of Sciences, Wuhan, China; University of Chinese Academy of Sciences, Beijing, China

**Keywords:** mutation prediction, generative deep learning, protein language model, *in silico* virus evolution

## Abstract

Predicting the mutation prevalence trends of emerging viruses in the real world is an efficient means to update vaccines or drugs in advance. It is crucial to develop a computational method for the prediction of real-world prevalent SARS-CoV-2 mutations considering the impact of multiple selective pressures within and between hosts. Here, a deep-learning generative framework for real-world prevalent SARS-CoV-2 mutation prediction, named ViralForesight, is developed on top of protein language models and *in silico* virus evolution. Through the paradigm of host-to-herd *in silico* virus evolution, ViralForesight reproduced previous real-world prevalent SARS-CoV-2 mutations for multiple lineages with superior performance. More importantly, ViralForesight correctly predicted the future prevalent mutations that dominate the COVID-19 pandemic in the real world more than half a year in advance with *in vitro* experimental validation. Overall, ViralForesight demonstrates a proactive approach to the prevention of emerging viral infections, accelerating the process of discovering future prevalent mutations with the power of generative deep learning.

## 1 Introduction

The representative emerging viral infection COVID-19, caused by Severe acute respiratory syndrome coronavirus 2 (SARS-CoV-2) [1, 2], has had a catastrophic impact on public health worldwide, including extremely high morbidity and millions of deaths ^1^. Regardless of geographic location, SARS-CoV-2 evolves within and between hosts [3], involving the action of multiple selective pressures [4–7]. The evolutionary trend of SARS-CoV-2 is profoundly influenced by those selective pressures [8], which causes its adaptive evolution to produce various variants, most notably multiple sublineages of Omicron [9].

Predicting the mutation prevalence trends of emerging viruses is a powerful means to update vaccines or drugs in advance. However, the significant experimental cost of deep mutational scanning (DMS) profiles [8, 10–19] makes it difficult for them to perform real-time predictive analysis. Existing computational methods for virus prediction also have limitations. First, the property prediction models of viral proteins [20–28] were customized according to specific properties, thus lacking a comprehensive consideration of selective pressures. Second, the methods of reproducing evolutionary trajectories [29, 30] can only perform post hoc analyses of the observed variants, and cannot predict the prevalence of unknown lineages. Third, the evolutionary trend prediction models [22, 31–33] did not consider both intra-host and inter-host selective pressures simultaneously. Therefore, it is crucial to develop a computational method for predicting prevalent mutations that spans from the host level to the herd level.

As shown in Fig.1a, as the link between intra-host evolution and inter-host evolution of SARS-CoV-2, the transmission bottleneck is often very narrow, in which variants with significant selective advantages are more likely to infect new hosts in a single transmission event [3, 34]. With continuous spreading, variants with significant selective advantages account for most of the transmission chains, thus showing the dominance of the pandemic at the herd level [3]. In order to make the computational methods take into account the selective pressures of intra-host evolution and inter-host evolution at the same time, it is a feasible strategy to introduce herd-level mutational drivers on top of the host-level mutational drivers. As shown in Fig.1b, for the intra-host adaptive evolution of SARS-CoV-2, ACE2 binding affinity, expression, and antibody escape are the main mutational drivers, i.e., host-level selective pressures [12, 35, 36]. First, according to a large number of previous studies [37–46], ACE2 binding affinity can be partially sacrificed during the intrapandemic evolution of SARS-CoV-2. Therefore, we do not consider the ACE2 binding affinity of variants at the host level in this work to avoid filtering out high-risk variants with reduced ACE2 binding affinity. Second, considering the importance of breaking through antibody immunity at the herd level for the occurrence of transmission events, we upgrade host-level antibody escape with estimated herd-level immunity barrier [47, 48].

**Fig. 1.**
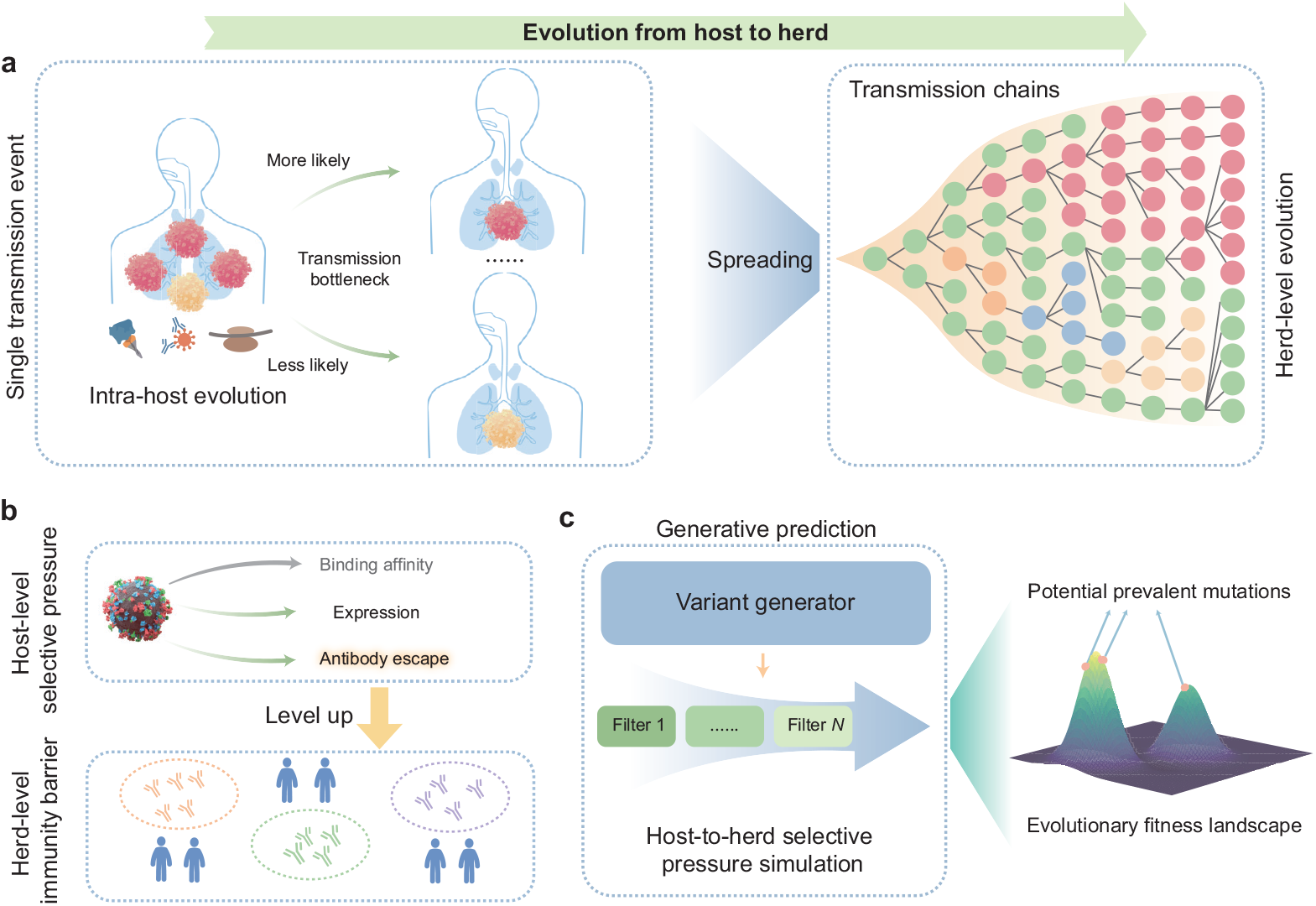
Our motivation and methodology. **a**, Illustration of the SARS-CoV-2 evolution from the host level to the herd level. After undergoing intra-host evolution, the variants spreads through the transmission bottleneck in a single transmission event (left panel), thus forming a large number of transmission chains, in which lineages with selective advantages become dominant (right panel). **b**, Our host-to-herd selective pressure simulation strategy. Intra-host adaptive evolution of SARS-CoV-2 involves main host-level selective pressures including ACE2 binding affinity, expression, and antibody escape, among which host-level antibody escape is upgraded to herd-level immunity barrier to integrate herd-level selective pressures of SARS-CoV-2. **c**, The methodology of our deep-learning generative prediction framework for potential prevalent SARS-CoV-2 mutations. Massive variants generated by the variant generator are subjected to host-to-herd selective pressure simulation, thereby recommending potential real-world prevalent mutations in the future in the vast evolutionary fitness landscape. **Alt text**: A three-panel schematic illustrating the motivation and methodology.

With the above host-to-herd selective pressure simulation strategy, a deep-learning generative framework for real-world prevalent SARS-CoV-2 mutation prediction, named ViralForesight, is developed on top of protein language models and *in silico* virus evolution. ViralForesight performs generative prediction through host-to-herd *in silico* virus evolution, including variant generation, selective pressure screening, and mutation ranking. The protein language models fine-tuned by real-time evolutionary information are employed as the variant generators for mutated-site-guided variant generation, and the predictive models of intra-host and inter-host selective pressures are adopted as diverse filters for variant screening (Fig.1c). Subsequently, the predicted SARS-CoV-2 mutations are ranked and recommended in the vast evolutionary fitness landscape, not only reproducing previous real-world prevalent mutations with superior performance, but also correctly predicting future prevalent mutations that dominate the COVID-19 pandemic in the real world more than half a year in advance with *in vitro* experimental validation. Overall, ViralForesight facilitates the predictive discovery of future prevalent mutations and provides insights for proactive prevention of emerging viral infections.

## 2 Results

### 2.1 A generative prediction framework across host to herd

ViralForesight is a generative prediction framework across host to herd to predict potential prevalent SARS-CoV-2 mutations in the future. Four modules make up this deep-learning framework, namely protein language model (PLM) fine-tuning, mutated-site-guided variant generation, host-to-herd selective pressure screening, and mutation ranking. Following this host-to-herd *in silico* virus evolution process, the top-ranked point mutations are recommended as the most likely prevalent SARS-CoV-2 mutations in the real world.

First, for the PLM fine-tuning module, the real-time SARS-CoV-2 evolutionary information is adopted to fine-tune the pretrained PLM to internalize evolutionary patterns of SARS-CoV-2 proteins (Fig.2a). Specifically, considering the ability of PLM to internalize evolutionary patterns [49–52], we choose the pretrained PLM (ESM-2 with 650M parameters here) as the basic variant generator. However, the off-the-shelf PLM is pretrained on massive natural protein sequences and lacks understanding of the mutational patterns of SARS-CoV-2 proteins. Therefore, we integrate the variant sequences of specific SARS-CoV-2 lineages to fine-tune the pretrained PLM to adjust the learned evolutionary patterns towards the mutational fitness of SARS-CoV-2 proteins (Supplementary information S3).

**Fig. 2.**
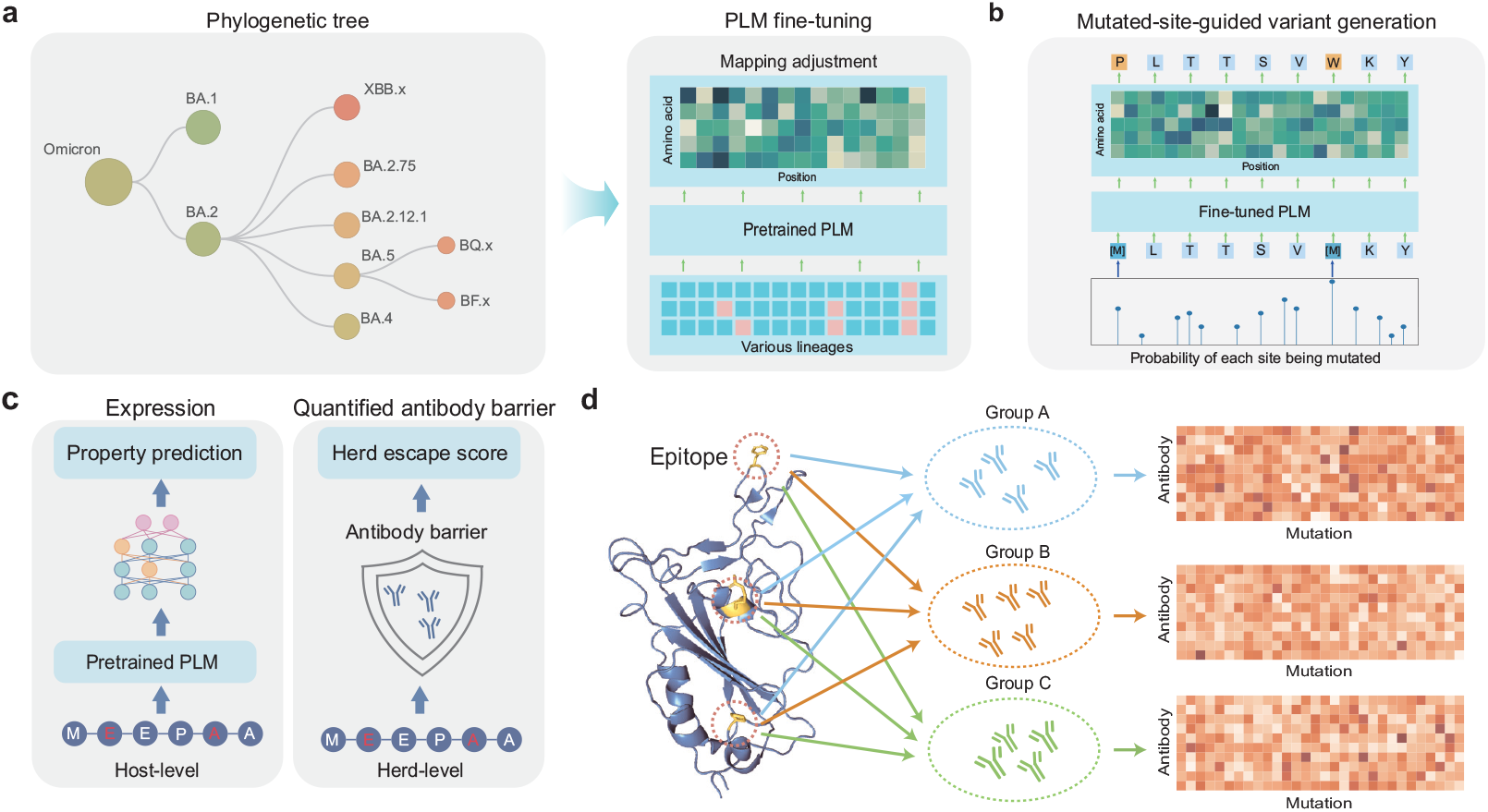
Module details. **a**, Protein language model (PLM) fine-tuning module, in which variant sequences of specific SARS-CoV-2 lineages carrying real-time evolutionary information are adopted to fine-tune the pretrained PLM. **b**, Mutated-site-guided variant generation module, where the probability of each site being mutated in real-time SARS-CoV-2 evolutionary trajectory is used as the probability of each site being masked (where to mutate) for the prediction of residue type to be mutated to (how to mutate). **c**, Host-to-herd selective pressure screening module, in which the *in silico* generated variant sequences are screened through the selective pressures of SARS-CoV-2 at host-level (expression prediction model) and herd-level (quantified antibody barrier model). **d**, Illustration of the proposed quantified antibody barrier model. Based on the deep mutational scanning data, monoclonal antibodies isolated from COVID-19 convalescent individuals are divided into multiple groups according to antigenic epitope, and the herd escape score of a variant is obtained by the average escape score of each of its mutations for each group. **Alt text**: A four-panel schematic detailing the modules of the proposed framework.

Second, after PLM fine-tuning, the mutated-site-guided variant generation module is conducted to simulate the real-world mutation process of SARS-CoV-2 RBD (Fig.2b). In general, the mutation process is determined by two factors, i.e. where to mutate (determine the site to be mutated) and how to mutate (determine the mutation type at that site). For the determination of mutated sites, the site-by-site mutated probabilities of real-time evolutionary trajectory of the starting lineage are adopted to determine the sites to be mutated during variant generation. Specifically, we calculate the distribution of mutated probabilities of all sites in the integrated variant sequence set for PLM fine-tuning that carries the real-time evolutionary trajectory information. Subsequently, taking the starting lineage as the reference sequence of *in silico* evolution, the probability of each site being mutated is used as its masked probability, determining where to mutate. Finally, the fine-tuned PLM predicts the residue type to be mutated to, determining how to mutate. In this way, massive variants are generated on demand.

Third, the host-to-herd selective pressure screening module is designed to screen the *in silico* generated variants through the selective pressures of SARS-CoV-2 proteins in two levels, i.e. host-level that determines survival and herd-level that determines transmission (Fig.2c). Based on our host-to-herd selective pressure simulation strategy, we developed a host-level expression prediction model (Fig.2c left panel) and a quantified antibody barrier model to quantify the capability of variants to break through the herd-level immune barrier with herd escape scores (Fig.2c right panel). More details can be found in Methods 4.2 and Supplementary information S5.

Fourth, the mutation ranking module is conducted to recommend potential prevalent SARS-CoV-2 mutations in the future. Specifically, we first retain the variants with enhanced RBD expression (classification probability larger than 0.5) at the host level, and then further retain the top 50% variants in the herd escape score ranking list. Subsequently, we calculate the frequency of each mutation in the screened variant sequences. Finally, mutations are ranked from high frequency to low frequency, where the top-ranked mutations are considered to have the potential to become prevalent in the future.

### 2.2 Prediction reliability of ViralForesight

To ensure the reliability of ViralForesight, we conducted multiple ablation experiments from three perspectives, including the completeness of generation scale, prediction stability, and screening effectiveness. In the retrospective experimental scenario of this section, we defined a SARS-CoV-2 lineage as starting lineage (the reference sequence of *in silico* evolution), and a lineage as cutoff lineage (the endpoint of variant sequence collection for PLM fine-tuning). Variant sequences between the starting and cutoff lineages were integrated into a sequence set for the PLM fine-tuning and calculation of mutated probabilities of all sites. The real-world prevalent mutations of the lineages emerging after the cutoff lineage along the phylogenetic tree were target mutations, i.e., the ground truth.

First, to avoid missing potential prevalent mutations with strong immune escape ability across the population, we performed ablation experiments for variant generation scale to determine a reasonable generation scale that covered the upper limit of fitness of *in silico* SARS-CoV-2 evolution. In those ablation experiments, BA.2.1 was adopted as the starting lineage, with BA.3 as the cutoff lineage. Specifically, after fine-tuning the PLM using the integrated variant sequence set from BA.2.1 to BA.3 lineage, variant generation was conducted at different scales (Supplementary information S7). As shown in Fig.3a, for both types of quantified antibody barrier models, the generation scale of one million made the escape capability increment begin to stabilize and approach 0 when setting different *K*, which appeared to reach the upper limit of herd-level immune escape ability. In addition, we conducted ablation experiments for binding affinity increment and expression increment to ensure the reasonableness of the generation scale (Supplementary information Fig.S1). Therefore, we set the variant generation scale to one million in all experiments of this work.

**Fig. 3.**
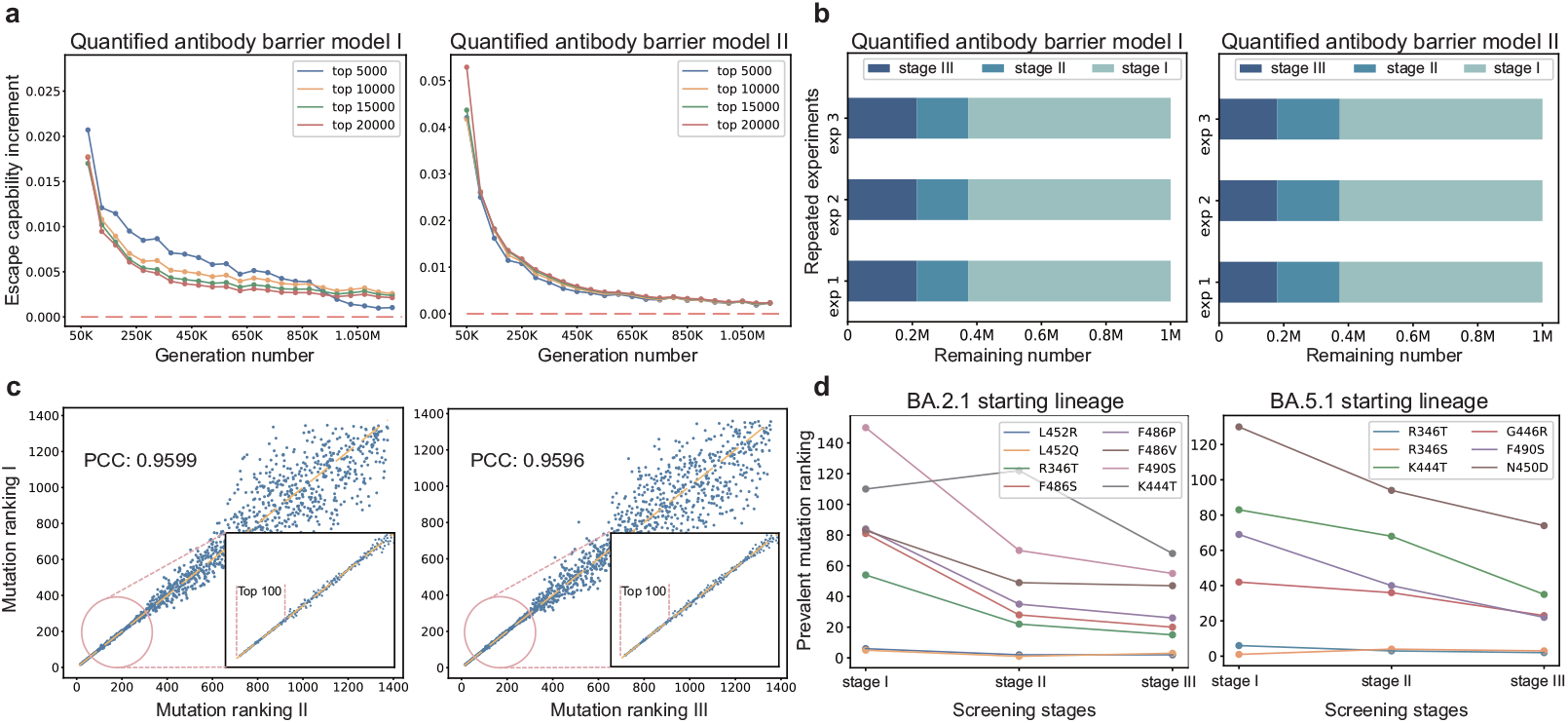
Ablation experiments. **a**, Ablation experiments for variant generation scale under two types of quantified antibody barrier models. The x-axis represents the number of generated variants with an interval of 50,000 and the y-axis represents the escape capability increment referring to the difference in the average herd escape score of the top *K* (*K* = 5, 000, 10, 000, 15, 000, 20, 000) variants sorted by scores in two consecutive generation experiments. **b**, The remaining number of variants at different screening stages of three repeated experiments under two types of quantified antibody barrier models. “Stage I” refers to the stage of initial variant generation, “Stage II” refers to the stage where the variants are screened by the expression prediction model, and “Stage III” refers to the stage where the variants are further screened by the quantified antibody barrier model. **c**, The Pearson correlation coefficient (PCC) in mutation rankings across three repeated experiments. **d**, The ranking trends of previous real-world prevalent mutations (target mutations) for BA.2.1 (left panel) and BA.5.1 (right panel) across different screening stages. **Alt text**: A four-panel figure presenting ablation experiments.

Second, considering the PLM fine-tuning and the variant generation modules may involve randomness to the whole prediction pipeline, we conducted experiments from two perspectives to test the prediction stability of ViralForesight, including quantitative consistency across different screening stages (Fig.3b) and correlation of the mutation rankings in repeated experiments (Fig.3c). In those ablation experiments, BA.2.1 was adopted as the starting lineage, with BA.3 as the cutoff lineage. On the one hand, as shown in Fig.3b, we conducted three repeated experiments under the two types of quantified antibody barrier models to observe the remaining number of variants at different screening stages. It can be noticed that regardless of the type of quantified antibody barrier model, the number of variants in each screening stage was very close (Supplementary information Table S1 and Table S2) in the three repeated experiments, demonstrating strong stability from the perspective of quantitative consistency. On the other hand, as shown in Fig.3c, the correlation in mutation rankings was observed across three repeated experiments. We can find that the mutation rankings of the three repeated experiments showed strong correlation, with the Pearson correlation coefficient (PCC) exceeding 0.95. It is worth noting that the higher the ranking of the mutation, the better the correlation in the repeated experiments, and the top 100 mutations were close to perfect correlation, which ensured that our prediction results were stable because we ultimately recommended the top-ranked mutations as candidates for prevalent mutations in the future.

Third, to demonstrate the effectiveness of selective pressure screening, we visualized the ranking trends of previous real-world prevalent mutations (target mutations) across different screening stages in Fig.3d. For those ablation experiments, in addition to BA.2.1 as the starting lineage, BA.5.1 was also adopted as the starting lineage, which was paired with BQ.1 as the cutoff lineage. When BA.2.1 was the starting lineage, there were 14 target mutations (339H, 346T, 368I, 444T, 445P, 446S, 452Q, 452R, 460K, 478R, 486P, 486S, 486V, 490S), while when BA.5.1 was the starting lineage, there were 7 target mutations (346S, 346T, 444T, 446R, 450D, 460K, 490S). We can find that 8 of the 14 target mutations in BA.2.1 (i.e., 57%, Fig.3d left panel) and 6 of the 7 target mutations in BA.5.1 (i.e., 86%, Fig.3d right panel) continued to rank higher with the implementation of selective pressure screening models, including the expression prediction model in Stage II and two types of quantified antibody barrier models in Stage III. Overall, most of the previous real-world prevalent mutations were ranked higher and higher as the screening models were gradually executed, proving the contribution of the host-to-herd selective pressure screening module to the recommendation of potential prevalent mutations.

Fourth, we conducted ablation experiments to explore the impacts of PLM finetuning and mutated-site-guided generation on the prediction results in Supplementary information S10. In addition, the impact of introducing the binding affinity screening model into the ViralForesight framework was also quantified in Supplementary information S12. More importantly, we also performed experiments with B.1.1 as the starting lineage for the evolution prediction of Omicron (B.1.1.529) in Supplementary information S13. The above results further demonstrated the prediction reliability and broader applicability of ViralForesight.

### 2.3 Reproduction of previous real-world prevalent mutations

To demonstrate the predictive power of ViralForesight, we performed *in silico* validation experiments on the recurrence of previous prevalent mutations taking BA.2.1 and BA.5.1 as the starting lineage, respectively. The settings of the *in silico* validation experiments were consistent with those of ablation experiments in Section 2.2. In addition, a certain previous prevalent mutation ranked within the top 100 by ViralForesight was considered to be correctly predicted.

The state-of-the-art method for predicting specific mutations through *in silico* directed evolution, MLAEP [31], was used to compare the prediction performance with our ViralForesight. Considering that MLAEP, which is based on genetic algorithms, relies heavily on the diversity of starting sequences, we included lineages before Omicron for better performance when implementing MLAEP, while only lineages after Omicron were included for ViralForesight. In general, the number of previous prevalent mutations correctly predicted by ViralForesight (i.e., the prevalent mutations ranked within top 100) was two to seven times that of MLAEP (11 vs. 5 for BA.2.1 and 7 vs. 1 for BA.5.1), and all the correctly predicted prevalent mutations of MLAEP were included in the predicted ranking list of ViralForesight. In other words, even though MLAEP used more starting lineages, ViralForesight’s hit rate for previous real-world prevalent mutations was still 42.9% higher than that of MLAEP in BA.2.1 lineage (78.6% vs. 35.7%) and 85.7% higher in BA.5.1 lineage (100% vs. 14.3%), respectively. We also provided the prediction results of MLAEP with only Omicron lineages as starting sequences in Supplementary information Fig.S2, and the diversity of the predicted mutation sites decreased significantly.

As shown in Fig.4ab, the prediction performance of our ViralForesight and MLAEP is presented in the form of sequence logos [53, 54], where sites carrying previous real-world prevalent mutations are selected to visualize the distribution of predicted mutations. The heights of the letters corresponding to the correctly predicted previous real-world prevalent mutations in ViralForesight’s sequence logos were more uniform than those in MLAEP, indicating that ViralForesight ranked the ground-truth target mutations higher overall. To show the improvement of ViralForesight’s predicted ranking for previous prevalent mutations in more detail, we compared the rankings of correctly predicted target mutations in ViralForesight and MLAEP in Fig.4c, where mutations ranked after 100 are represented by “>100” for better visualization. We can find that, compared with MLAEP, our ViralForesight gave significantly higher rankings to previous real-world prevalent mutations, demonstrating a significant improvement in prediction performance.

**Fig. 4.**
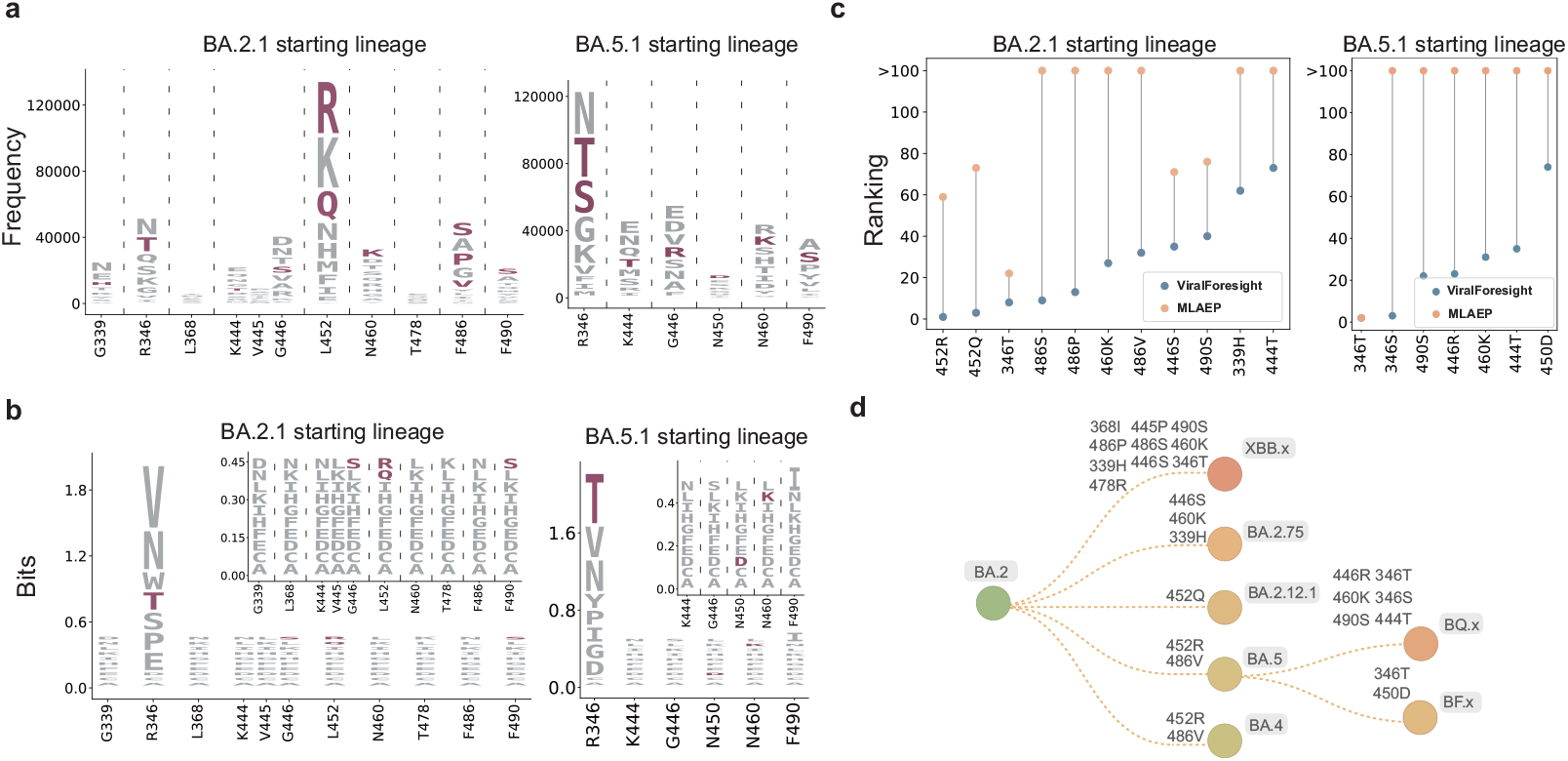
Reproduction of previous real-world prevalent SARS-CoV-2 mutations. **ab**, Sequence logos of the predicted mutations of our ViralForesight (a) and the state-of-the-art method MLAEP (b) on the sites carrying previous real-world prevalent mutations with BA.2.1 (left panel) or BA.5.1 (right panel) as the starting lineage. A certain previous prevalent mutation ranked within the top 100 by ViralForesight or MLAEP is considered to be correctly predicted and colored in purple. For MLAEP, the sequence logos at sites other than R346 are enlarged for clear observation. **c**, Ranking improvement of correctly predicted previous real-world prevalent mutations with BA.2.1 (left panel) or BA.5.1 (right panel) as the starting lineage, in which the mutations ranked after 100 are uniformly represented by “*>*100”. **d**, Annotated phylogenetic tree, where previous real-world prevalent mutations correctly predicted by our ViralForesight are annotated on the corresponding lineages. The “x” suffix represents a collection of related lineages. **Alt text**: A four-panel figure comparing predicted SARS-CoV-2 mutations of different methods.

As shown in Fig.4d, we annotate the previous real-world prevalent mutations correctly predicted by ViralForesight on the corresponding lineages in the phylogenetic tree. We can find that whether BA.2.1 or BA.5.1 was used as the starting lineage, there were correctly predicted real-world prevalent mutations on the subsequent lineages. When BA.2.1 was taken as the starting lineage, although XBB lineages appeared relatively late among its subsequent lineages, the multiple prevalent mutations they contain were correctly predicted (the top panel of Fig.4d). This indicates that the potential prevalent mutations predicted by ViralForesight may not appear in the near future, but are likely to spread at a certain time point, which is closely related to the element of chance introduced by the transmission bottlenecks and geographical locations [3, 34, 55].

### 2.4 Recommendation of future prevalent mutations more than half a year in advance

After confirming ViralForesight’s ability to reproduce previous prevalent mutations, we explored how to use it to recommend potential prevalent mutations in the future. Specifically, the predictions of ViralForesight were validated *in vitro* to recommend mutations with potential to become prevalent in the future, with XBB.1.5 as the starting lineage. Unlike the retrospective experiments in the previous sections, the predictive experiments in this section do not involve a cutoff lineage. The recommendation pipeline consists of 5 steps as described in Supplementary information S14.

Entry efficiency is indicative of the viral invasiveness into host cells [56, 57], and viral evasion of neutralizing antibodies could exacerbate disease symptoms and facilitate transmissions [58–60]. Therefore, as shown in Fig.5ab, for the top 12 point mutations predicted by ViralForesight, we performed entry efficiency and neutralization experiments to determine their influences on viral entry efficiency and immune evasion.

**Fig. 5.**
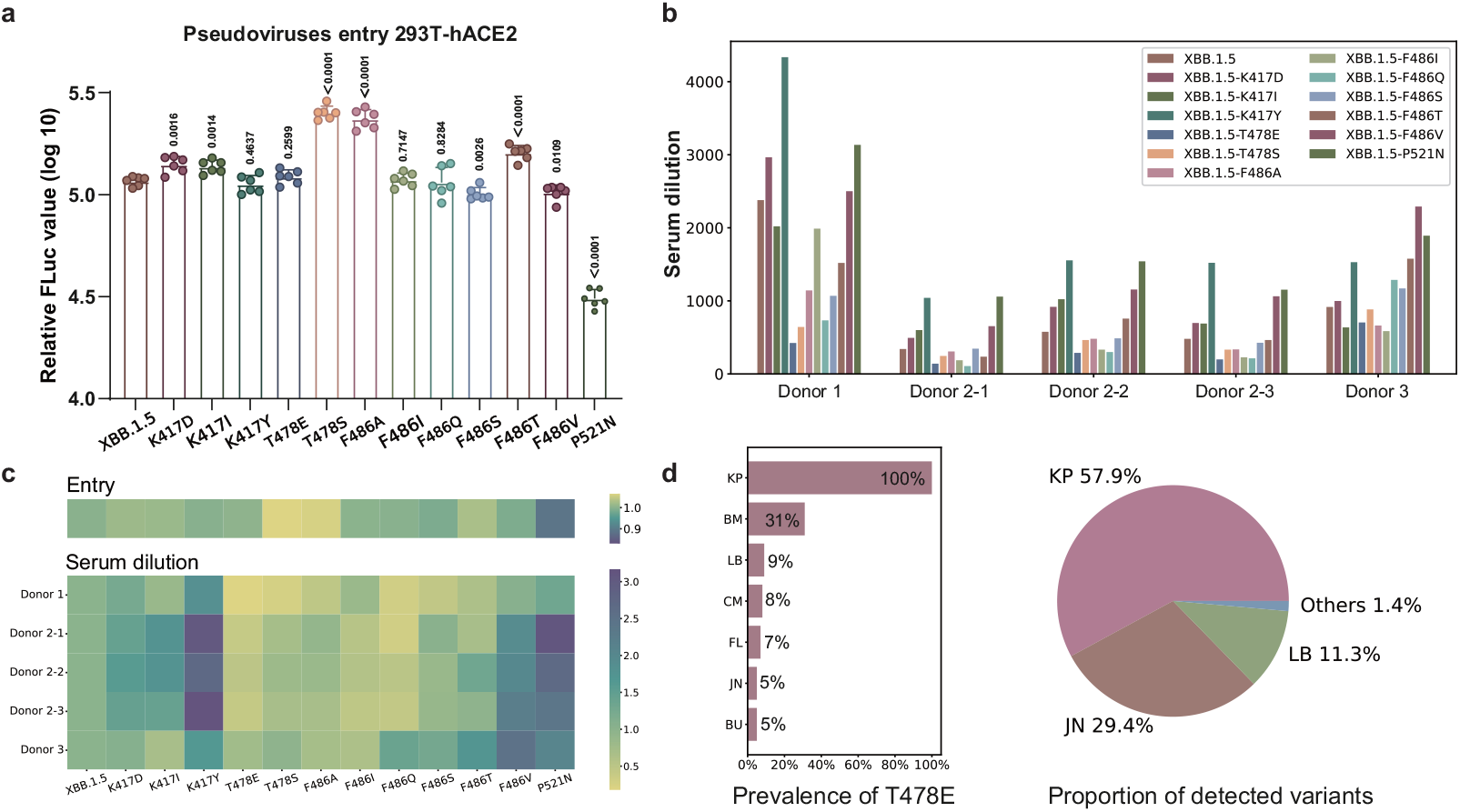
*In vitro* validation and real-world prevalence analysis of recommended SARS-CoV-2 mutations. **a**, VSV-FLuc-based pseudotyped virus entry. Error bars revealed the standard deviation of the means from six biological repeats and the data were analyzed by Student’s t-test. **b**, Neutralizing assay with convalescent sera. Neutralizing IC50 of sera samples to different spike-pseudotyped viruses was calculated by measuring the FLuc activity. **c**, Summary of entry efficiency and serum dilution. Entry efficiency (upper panel) and serum dilution to reach IC50 (lower panel) of predicted mutated spike pseudoviruses are compared with original XBB.1.5 spike pseudovirus. For entry assay, yellow color means higher viral infectivity, and for neutralizing assay, yellow color means stronger immune evasion. **d**, Real-world prevalence analysis of recommended representative T478E mutation, including the mutation prevalence of T478E from outbreak.info and GISAID at June 2024 (left panel) and the proportion of detected variants from CDC COVID Data Tracker at the collection week of June 24, 2024 (right panel). **Alt text**: A four-panel figure showing *in vitro* validation and real-world analysis of predicted SARS-CoV-2 mutations.

First, in order to investigate the impact of the top 12 single-amino-acid mutations on viral entry stage in susceptible cells, vesicular stomatitis virus (VSV)-derived pseudovirus assays were conducted. As shown in Fig.5a, compared with the original XBB.1.5 spike protein, seven variants with single-amino-acid mutations, including K417D, K417I, T478E, T478S, F486A, F486I, and F486T, exhibited higher FLuc activity. Notably, T478S, F486A, and F486T showed significantly increased entry efficiency (P value < 0.0001), suggesting enhanced invasiveness into susceptible cells. Conversely, mutations like F486S, F486V, and P521N in the RBD of XBB.1.5 reduced infection in 293T-hACE2 cells, particularly the P521N mutation. Mutations such as K417Y, T478E, F486I, and F486Q did not affect the entry process significantly, with minimal differences in entry efficiency compared to the original XBB.1.5 spike protein.

Second, the evasive capabilities of these variants carrying predicted mutations for neutralizing antibodies were further evaluated. As shown in Fig.5b, among the predicted mutations, T478E, T478S, F486A, and F486I demonstrated substantial immune evasion against all five sera tested. F486Q and F486S attenuated the neutralizing ability of four sera, whereas F486T was not effectively neutralized by three sera.

According to the overall results of the *in vitro* validation in Fig.5c, T478E, T478S, F486A, and F486I appeared to pose greater risks among the 12 predicted mutations, as they displayed increased entry efficiency in susceptible cell lines and enhanced immune evasion against all the convalescent sera compared with the original XBB.1.5 spike protein. Therefore, the above four mutations were recommended for real-world prevalence analysis. Specifically, with GISAID ^2^, outbreak.info ^3^, and CDC COVID Data Tracker ^4^ as references, the above four mutations were tracked for prevalence in the real world at June 2024. According to the prevalence data of real world, three of the four mutations (namely T478E, F486A, F486I) had appeared at extremely high frequency (more than 80% of all sequences) in multiple lineages during the period from the time of *in vitro* experimental validation (December 2023) to the time of prevalence analysis (June 2024), confirming their widespread prevalence in the real world.

As shown in Fig.5d left panel, taking the representative mutation T478E as an example, its mutation prevalence from outbreak.info and GISAID at June 2024 showed that 100% of the variant sequences with the prefix KP contained this mutation, and the prefixes of the remaining lineages containing this mutation that account for no less than 5% were BM (31%), LB (9%), CM (8%), FL (7%), JN (5%), and BU (5%), respectively. Meanwhile, as shown in Fig.5d right panel, the proportion of detected variants from CDC COVID Data Tracker (the collection week of June 24, 2024) showed that the lineage prefixed with KP was dominant (57.9%), and the remaining lineages with a proportion of more than 10% were prefixed with JN (29.4%) and LB (11.3%). Combining the above results, we can find that the prevalence of predicted T478E had a clear correlation with the proportion of variants detected in the real world, which indicated that the point mutations recommended by our prediction pipeline not only occurred widely in the real world, but also had the potential to dominate the pandemic.

## 3 Discussion

As a generative deep-learning framework, ViralForesight achieves prediction of potential prevalent SARS-CoV-2 mutations in a huge evolutionary space through *in silico* virus evolution based on the host-to-herd selective pressure simulation strategy. ViralForesight still has limitations, including supporting only prediction of single-site mutations (residue substitutions) and the lack of integration of high-level complex factors in the model. First, the lack of systematic mutation data for multi-site mutations, residue insertions and deletions in SARS-CoV-2 makes it difficult to train a ViralForesight that supports the mutation prediction for multi-site or non-equal length. Second, the high-level complex factors involved in viral evolution, including transmissibility, pathogenicity, virulence, and geographic location, lack refined machine learning modeling strategies, limited by insufficient data or complex dynamic changes. For transmissibility, it is gratifying that there are some mathematical forms for SARS-CoV-2 infectiousness [61, 62], but the difficulty of obtaining real-time transmission data cannot be ignored. For pathogenicity and virulence, the isolation of existing clinical data between different hospitals makes it difficult to integrate for machine learning modeling. For geographic location, the different public health strategies adopted by different countries or regions make the impact of geographic location complex and variable, significantly increasing the difficulty of using mathematical forms to quantitatively describe it. Integrating these high-level complex factors is necessary because geographic and demographic factors can indeed influence viral evolution by shaping transmission dynamics, selective pressures, and genetic drift. For instance, high-population-density regions may facilitate the selection of immune escape mutations due to increased transmission rates, while geographic isolation could lead to region-specific lineage diversification. Therefore, in the future, introducing some reasonable mathematical forms for transmissibility, developing generalizable machine learning models based on a small amount of clinical data for pathophysiology, and modeling the impact of geographic location as finely as possible are potential directions to further improve the predictive performance of ViralForesight.

Last but not least, we acknowledge that while such mutation predictions have great potential for preparedness, they must be handled responsibly to prevent misuse and unintended consequences. By adhering to established biosecurity guidelines and collaborating with policymakers, we can ensure that predictive models serve as tools for preparedness rather than sources of unintended risk (Supplementary information S16).

## 4 Methods

### 4.1 Datasets

The sequence set for PLM fine-tuning and variant generation were integrated from the GISAID database ^5^ at December 2023 (Login/EpiCoV/Downloads/Alignment and proteins). For host-level expression prediction, the deep mutational scanning (DMS) datasets of previous work [12] were adopted to train a predictive model. We only used the samples of Omicron BA.1 and Omicron BA.2 due to the notable difference in lineages before and after Omicron [63, 64]. For herd-level escape score prediction, two sets of DMS datasets from previous studies [8, 13] were adopted to build the antibody barrier models that quantified the capability of variants to break through the herd-level immune barrier. Since Omicron sub-lineages were prevalent during the time of this work, we only used the DMS data of antibodies isolated from convalescents of Omicron variants. The details of dataset selection and preprocessing can be seen in Supplementary information S1.

### 4.2 Screening models

The host-level expression prediction model adopted the deep-learning framework proposed by a previous work [33], which included three steps, namely sequence embedding extraction, local-global dependence coupling and multi-task focal learning. The herd-level quantified antibody barrier model was built on the DMS datasets from the previous studies [8, 13]. For each variant sequence, it contains many mutations, such as G339H, R346T, etc. The mutations on different positions would bring different effects on the herd-level antibody escape. Therefore, we first divided different mutations into different groups according to antigenic epitope, and calculate the mutational effects in each group. Then we summed up the effects of different groups based on the group importance. The implementation details of the above screening models can be seen in Supplementary information S5.

### 4.3 Step-by-step experimental reproduction instructions

ESM-2 (650M) was adopted as the pretrained PLM for further finetuning and property prediction. For data preprocessing, the variant sequences from GISAID were filtered based on the Pango nomenclature [65]. The variants between starting lineage and cutoff lineage were taken for PLM finetuning with a masked residue modeling task (the same task with ESM-2 pretraining) for 1 epoch. The whole model was fine-tuned with AdamW optimizer, and the learning rate was set as 1 *×* 10^−5^. For variant generation, the number of generated sequences was set as 1 million. For expression prediction model settings, the CNN layer number, kernel size and stride were set as 3, 3 and 1 respectively. The hidden dimension of ContextPool-attention was 1,024. For prediction head, we used 4 layers of Linear layer, and the dropout rate is 0.6. Leaky ReLU was adopted as an activation function and the Adam optimizer was adopted for model training. The learning rate was 2 *×* 10^−5^, and the training epoch was 200. For mutation ranking, we set the expression threshold as 0.5 (i.e., the variants with output probabilities of classification prediction larger than 0.5 were retained), and retained the sequences with top 50% herd escape scores. For the remaining sequences, we calculated the frequency of each mutation, and the top 100 mutations were considered to have the potential to become prevalent in the future.

### 4.4 *In vitro* experimental validation

First, pseudoviruses containing original or mutated spike proteins of XBB.1.5 were packaged and diluted to equal copy numbers. Second, 293T-hACE2 cells were infected by pseudoviruses, and the firefly luciferase (FLuc) activity of infected cells was quantified. Third, Sera from three convalescent individuals were serially diluted and incubated with an equal volume of pseudoviruses at 37 °C. Fourth, following incubation, 293T-hACE2 cells were infected with the serum-pseudovirus mixture. Finally, the half-maximal inhibitory concentration (IC50) of these sera against the original XBB.1.5 spike protein and its variants was determined by measured FLuc activity. More details can be seen in Supplementary information S15.

## Key Points

- We propose a generative deep-learning framework, ViralForesight, for real-world prevalent SARS-CoV-2 mutation prediction on top of protein language models and *in silico* virus evolution.
- We develop a host-to-herd selective pressure simulation strategy to incorporate host-level and herd-level virus evolutionary drivers.
- ViralForesight reproduced previous real-world prevalent SARS-CoV-2 mutations for multiple lineages with superior performance.
- ViralForesight correctly predicted the future prevalent mutations that dominated the COVID-19 pandemic in the real world more than half a year in advance with *in vitro* experimental validation.
- ViralForesight demonstrates a proactive approach to the prevention of emerging viral infections, accelerating the process of discovering future prevalent mutations with the power of generative deep learning.

## Biographical note

Dr Chen’s laboratory is committed to promoting the intersection of state-of-the-art deep-learning technologies and life sciences, focusing on foundation models in the field of life sciences and their applications to proteins and small molecules.

## Author contributions

Zhiwei Nie, Xudong Liu, and Haorui Si conceived the concept; Xudong Liu, Zhiwei Nie, and Haorui Si designed the deep-learning framework, trained models, and performed computational experiments. Xurui Shen performed *in vitro* experimental validation. Yutian Liu, Xiansong Huang, Tianyi Dong, and Fan Xu collected the data involved in the experiments and provided helpful discussions. Jie Chen, Peng Zhou, and Zhixiang Ren supervised this research and provided helpful discussions. All authors contributed to the writing and revision of the manuscript.

## Data and code availability

The sequence set for PLM fine-tuning and variant generation is freely available at https://gisaid.org/. The deep mutational scanning datasets of screening models are publicly available from previous studies [8, 12, 13]. The data used in our experiments can be reached at https://github.com/Kevinatil/ViralForesight. Relevant code and models are available at https://github.com/Kevinatil/ViralForesight.

## Footnotes

1 https://data.who.int/dashboards/covid19

2 https://gisaid.org/

3 https://outbreak.info/

4 https://covid.cdc.gov/covid-data-tracker/#datatracker-home

5 https://gisaid.org/

